# The effect of dietary fish oil replacement by microalgae on the gilthead sea bream midgut bacterial microbiota

**DOI:** 10.1101/2024.01.24.576938

**Authors:** Stefanos Katsoulis-Dimitriou, Eleni Nikouli, Elli Zafeiria Gkalogianni, Ioannis T. Karapanagiotidis, Konstantinos A. Kormas

## Abstract

It is well known that the gut microbiome and its interaction with the host influence several important factors for fish health such as nutrition and metabolism. Diet is one of the main factors influencing the composition of the gut microbiome in reared fish. Microalgae, due to their high fatty acid content, appear to be a promising alternative for replacing fish oil in aquafeed. Thus, the aim of this work was to evaluate the effects of dietary microalgae blends as fish oil replacers on the midgut bacterial microbiota of the gilthead sea bream (*Sparus aurata*). The control diet (FO) contained only fish oil as source of lipids, EPA and DHA fatty acids, while three experimental diets were used where fish oil was replaced at 67% by one of the following microalgae biomass blends: *Microchloropsis gaditana* and *Isochrysis* sp. (*Tisochrysis lutea*) (MI), *Phaeodactylum tricornutum* and *Isochrysis* sp. (PI) and *Schizochytrium* sp. and *P. tricornutum* (SP). The midgut bacterial community composition of the experimental diets was altered compared to the control diet. There were 11 operational taxonomic units (OTUs) which were highly abundant in FO compared to the three experimental diets (FO, MI, SP) and two OTUs that were found in high abundance in both FO and the experimental diets in all comparisons (FO-MI, FO-PI, FO-SP). Most of the highly abundant OTUs in the experimental diets were unique to each experimental diet, with two OTUs being found in common between FO-MI and FO-PI. Additional evidence from the presumptive bacterial functional metabolic pathways suggested that the microalgae-based diets resulted in one over-expressed and one under-expressed pathway. The overexpressed pathway was related to the metabolism of fucose, a major constituent of the polysaccharide content of several microalgal species. Peptidoglycan biosynthesis was the under-expressed metabolic pathway. This suggests that a new gut microbiota profile was selected due to the microalgae inclusion in the provided diet. This study showed that, with the absence of mortality in fish, the gilthead sea bream gut microbiome can smoothly adapt its function according to the metabolic capacity of the dietary microalgae combinations that were used. The MI feed seems to promote several beneficial bacteria with potential probiotic abilities in the fish gut, belonging to the *Pseudoalteromonas, Pseudomonas, Bacillus* and *Rhodopseudomonas* genera.

## INTRODUCTION

In reared fish, the composition of the gut microbiome is mainly determined by host selection (Roeselers et al. 2011) and diet (Liu et al. 2016), while environmental and other stochastic factors seem to be more important for wild fish (Kim et al. 2021, Kormas et al. 2023). The gut microbiome and its interactions with the host influence several important factors for fish health such as nutrition and metabolism (Liu et al. 2021), maintaining the development and maturation of the immune system (Koch et al. 2018), the nervous system functioning and fish development (Phelps et al. 2017). Furthermore, commensal bacteria have the potential to protect the fish against pathogens (Kelly & Salinas 2017) and when a disease occurs there is a loss of gut microbiota diversity (Felix et al. 2022). Gut microbiota dysbiosis is common in aquaculture species and is associated with changes in microbial structure, an increase in pathogenic microorganisms and/or decreases in the abundance of beneficial taxa (Xavier et al., 2023). For these reasons it is very important to investigate the changes that may occur in the composition of the gut microbiota with changes in the fish’s diet.

The main sources of proteins and lipids for carnivorous farmed fish, such as the gilthead sea bream (*Sparus aurata)*, are fish meals and fish oils, respectively, which are added in the aquafeeds in order to satisfy fish nutritional requirements and to provide high nutritional quality to the final product (Siddik et al. 2023). Fish meals and fish oils originate mainly from wild small pelagic fish species, making the aquaculture sector dependent on wild fish stocks and, thus, reducing their sustainability (Péron et al. 2010). As a result, several studies have focused on alternative sources of proteins and lipids to reduce the incorporation of fish meals and fish oils in aquafeeds (Oliva-Teles et al. 2015, Naylor et al. 2021). Among these alternatives, microalgae and oomycetes are suitable and sustainable feed ingredients.

Relevant studies showed that microalgae/oomycetes containing feeds have high digestibility, positive effect on the growth rate of aquatic species, high quantities of important pigments like carotenoids and phycobiliproteins, polysaccharides with potential antiviral and antibacterial properties, potential immunostimulant and probiotic effects and low carbon footprint for their production (Shah et al. 2017, Ahmad et al. 2022, Ma & Hu 2024).

*Microchloropsis gaditana* (Fawley, Jameson & Fawley 2015; formerly *Nannochloropsis gaditana*) is a marine microalga of the class Eustigmatophyceae with high content of eicosapentaenoic acid (EPA) (usually 36% of total fatty acids) (Zanella & Vianello 2020). *Phaeodactylum tricornutum* is a marine diatom of the class Bacillariophyceae, rich in EPA (29% of total fatty acids) (Matos et al. 2016). The oomycete genus *Schizochytrium* belongs to the Thraustochytriaceae family. Members of this genus are especially rich in docosahexaenoic acid (DHA) (41% of total fatty acids) (Karapanagiotidis et al. 2022). The marine microalga *Tisochrysis lutea* (here referred to as *Isochrysis* sp. due to its reclassification (Bendif et al. 2013) also produces high levels of DHA (49% of total fatty acids) (Ren et al. 2010).

Existing research has evaluated the use of *P. tricornutum* and *M. gaditana* in alternative feeds for the gilthead sea bream. In particular, *P. tricornutum* when used in experimental feeds, either alone or in combination with *Tetraselmis chui* and *Bacillus subtilis*, has been linked with reduced gut microbial diversity in gilthead sea bream (Cerezuela et. al 2012). However, the above combination along with dietary inulin and the utilization of its b-glycans, have demonstrated immunostimulant effects, influencing intestinal gene expression (Cerezuela et al. 2013; Reis et al. 2021; Teixeira et al. 2022). The use of *M. gaditana* showed that gut microbiota composition and intestinal integrity remained unaltered (Cerezo-Ortega et al. 2021) or an increase in intestinal microbiota richness without affecting intestinal morphology and function (Jorge et al. 2019). When used in combination with other species (*Chlorella* sp., *Arthrospira* sp., *Tisochrysis lutea, Scenedesmus almeriensis*) it was shown to favour the presence of probiotic bacteria in the gut and reduce the presence of pathogenic species (Garcia-Márquez et al. 2023; Garcia-Márquez et al. 2023). To the best of our knowledge there are no studies about the effects of *Isochrysis* sp. and *Schizochytrium* sp. on the gilthead sea bream gut microbiota.

These microalgal species when combined in fish diets, appear suitable for replacing the levels of EPA and DHA typically provided by fish oils. To our knowledge, there are no studies evaluating the potential differences on the midgut microbial composition and function of gilthead seabream when fed with a diet containing these microalgae species in combination. Thus, the aim of this study was to investigate the potential effect on the midgut microbial composition and function of gilthead seabream when fed a diet containing combination of *Phaeodactylum tricornutum, Isochrysis* sp., *Schizochytrium* sp. and *Microchloropsis gaditana* inclusions. It is hypothesised that microalgal-based diets will promote gut microbiota related to the metabolism of these microalgae.

## MATERIALS AND METHODS

### Sampling

For the needs of the present study, 40 *Sparus aurata* specimens (10 from each nutritional group, including the control diet) weighing 28-37g were collected from a feeding trial (Gkalogianni et al. 2023) conducted by FELASA accredited scientists at the experimental facilities of the Department of Agriculture, Ichthyology and Aquatic Environment registered for the use of experimental fish for scientific purposes (EL-43BIO/exp-01). Fish were obtained from a commercial hatchery (8 g of initial mean weight) and reared for totally 80 days into glass tanks within a closed seawater recirculation system (21.0±1.0 °C, pH at 8.0±0.2, salinity at 33±0.5 g/L, dissolved oxygen >6.5 mg/L, total ammonia nitrogen <0.1 mg/L. Fish were divided into 4 nutritional groups in triplicate tanks - with 25 fish each tank-with each group fed with a different diet. The four diets were isoproteic (48%), isolipidic (15.5%) and isoenergetic (21 Mj/kg) and satisfied the EPA+DHA requirements of the species. The control diet (FO) contained 80 g/Kg fish oil serving as a sole source of EPA and DHA, 40 g/Kg soybean oil and 250 g/Kg fishmeal with a EPA+DHA level at 2.8 g/Kg . Three other diets were formulated replacing 50% of the dietary fish oil level of the control diet by a blend of microalgae biomasses serving as sources of EPA and DHA: The PI diet contained 173 g/Kg *P. tricornutum* and 103 g/Kg *Isochrysis* sp. with EPA+DHA level at 2 g/Kg, the SP diet contained 39 g/Kg *Schizochytrium* sp. and 174 g/Kg *P. tricornutum* with EPA+DHA level at 2.6 g/Kg, and MI diet contained 137 g/Kg *M. gaditana* and 103 g/Kg *Isochrysis* sp. with EPA+DHA level at 1.9 g/Kg. Since the microalgae used in the experimental diets were in the form of dry biomass (and not extracted oils), their dietary inclusion contributed to the overall protein content. For this reason, the percentage of inclusion of fishmeal was replaced accordingly (at 44% in PI diet, 27% in SP diet and 36% in MI diet). Diets were pelletized at 1.5 mm diameter according to Karapanagiotidis et al. (2022). Fish promptly accepted the feeds and survival was high (99%) in all groups. At the end of the dietary trial, 10 individuals were randomly collected from each dietary group, euthanized with a strong dose of the anesthetic 2-phenoxyethanol (450 mg l^-1^, 15 min) and placed directly on ice. The body mass growth parameter W/L^3^ of the sampled fish based on weight (W) and length (L) ranged between 0.013±0.001 (FO) and 0.012±0.002 (MI) (Karapanagiotidis & Kormas unpubl. data). For the gut microbiota analysis, the whole intestine of each individual was removed in aseptic conditions, with scissors and scalpel after sterilization with 70% ethanol before each dissection and the midgut part was stored at -80 °C until the DNA extraction procedure.

### DNA extraction and sequencing

The DNA of seven gut samples of each feeding group (FO, PI, SP, MI) was extracted using the Qiagen DNeasy PowerSoil Pro Kit (Qiagen, Hilden, Germany), according to the manufacturer’s instructions. Bacterial communities were analysed by 16S rRNA gene amplicon sequencing targeting the V3-V4 hypervariable region with the primer pair S-D-Bact-0341-b-S-17 and S-D-Bact-0785-a-A-21 with Tm at 55°C (Klindworth et al. 2013). PCR amplification and sequencing was performed at MRDNA Ltd.1 (Shallowater, TX, United States) facilities on a MiSeq using paired end reads (2 × 300 bp) following the manufacturer’s guidelines.

### Data analysis

The 16S rRNA gene sequencing raw data were processed using the MOTHUR MiSeq standard of protocol procedure (Schloss et al. 2013); MOTHUR is a stand-alone bioinformatics platform covering the entire procedure from raw sequencing data to bacterial taxa 16S rRNA gene abundances. After trimming of the primer sequences ‘screen.seqs’ command was used with the parameters maxambig=0, minlength=331, maxhomop=8. The ‘split.abund’ command (cutoff = 1) was used to exclude the rare sequences. The operational taxonomic units (OTUs) at 97% cutoff similarity level were classified with the SILVA database release 138.1 (Quast et al. 2013, Yilmaz et al. 2014). The VSEARCH algorithm (Rognes et al. 2016) was used to detect and remove chimeric reads.

Sequences assigned as mitochondria or chloroplasts were removed from subsequent analyses. The ‘sub.sample’ command was used and the data were normalized to a depth of 2859 reads per sample maintaining, thus, a sufficient number of reads per sample but at the same time excluding only two samples with ≤2000 reads. Raw sequence reads have been deposited in the Short Read Archive (https://www.ncbi.nlm.nih.gov/sra) under the BioProject accession number PRJNA1068122. Identification of the closest relatives of the most abundant OTUs was performed using a Nucleotide Blast (http://blast.ncbi.nlm.nih.gov). The search was conducted with the following parameters: Standard databases, highly similar sequences (Megablast) and closest relatives were considered with percentage identity higher than 90%. The statistical analyses and graphic illustrations were performed using PAleontological STudies (PAST) software v.4.16 (Hammer et al. 2001). The input matrix for all statistical analyses was the OTUs table of each sequenced sample.

### Presumptive functional analysis of midgut microbiota

The PICRUSt2 v2.5.1 (Phylogenetic Investigation of Communities by Reconstruction of Unobserved STates) (Douglas et al. 2020) software was used to predict the microbiota functional profile (Langille et al. 2013). The predicted functions were searched in the MetaCyc database (Caspi et al. 2013), to be assigned to each metabolic pathway. Also, the KEGG (Kyoto Encyclopedia of Genes and Genomes) database (Kanehisa & Goto 2000, Kanehisa et al. 2023) was used to identify whether the genera of the most important OTUs had all the enzymes for specific metabolic pathways of interest.

## RESULTS

### Sequencing data and OTUs

After the quality filtering of the amplicon sequencing of the 16S rRNA gene V3–V4 region, a total of 319,103 reads remained. The number of reads ranged from 1,631 to 44,124 reads per sample. A subsample of 2,859 reads per sample was used for the normalization of the data. As a result, two samples (one from each of the FO and PI treatments) with only 1,631 and 1,867 reads, respectively, were excluded from the rest of the analyses. The total number of OTUs to which the reads were taxonomically assigned was 519.

### Alpha diversity

Taxa_S (OTUs richness), the Shannon_H and Simpson_1-D indices were calculated to assess the alpha diversity of the gut microbiota of gilthead sea bream in the four groups (Table 1). (Table 1). Compared to the control (FO), Shannon_H and Simpson_1-D values showed a slight increase in PI, while they decreased in MI and SP. Taxa_S increased in PI and SP and showed a slight decrease in MI. Despite the existing differences the permutational analysis of variance (PERMANOVA) calculated from the Bray-Curtis distance matrix performed with 9999 permutations (Table S1) showed no statistically significant differences between the indices values of the three experimental groups with FO. The only statistically significant difference was observed in Taxa_S (p=0.046) and Shannon_H (p=0.049) between PI and MI.

**Table 1.**
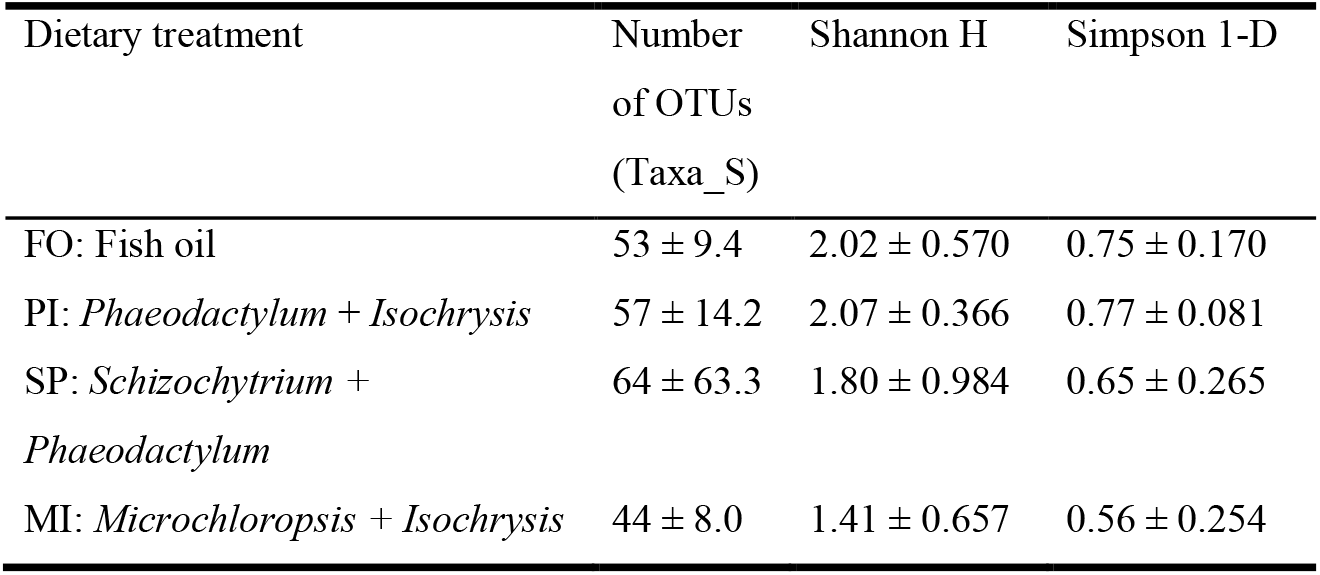
Alpha diversity metrics (Shannon H and Simpson 1-D), OTUs richness and Chao-1 values for each treatment. OTUs: operational taxonomic units.

### Beta diversity

Non-metric multidimensional scaling analysis (nMDS) was used to evaluate the beta diversity of the samples of the four feeding groups, based on a Bray-Curtis distance matrix (Figure 1). Samples from the MI groups showed the highest variation. The PI group was clearly separated from the samples of the FO and SP group. The samples of the FO, MI and SP groups overlapped and there was overlap between the samples of the PI and MI groups. Group PI was the only treatment with no overlap with FO. PERMANOVA pairwise test (Table S2) was used to further analyze the differences between dietary groups. Significant differences were observed between the PI and FO and between the PI and SP groups.

**Figure 1.**
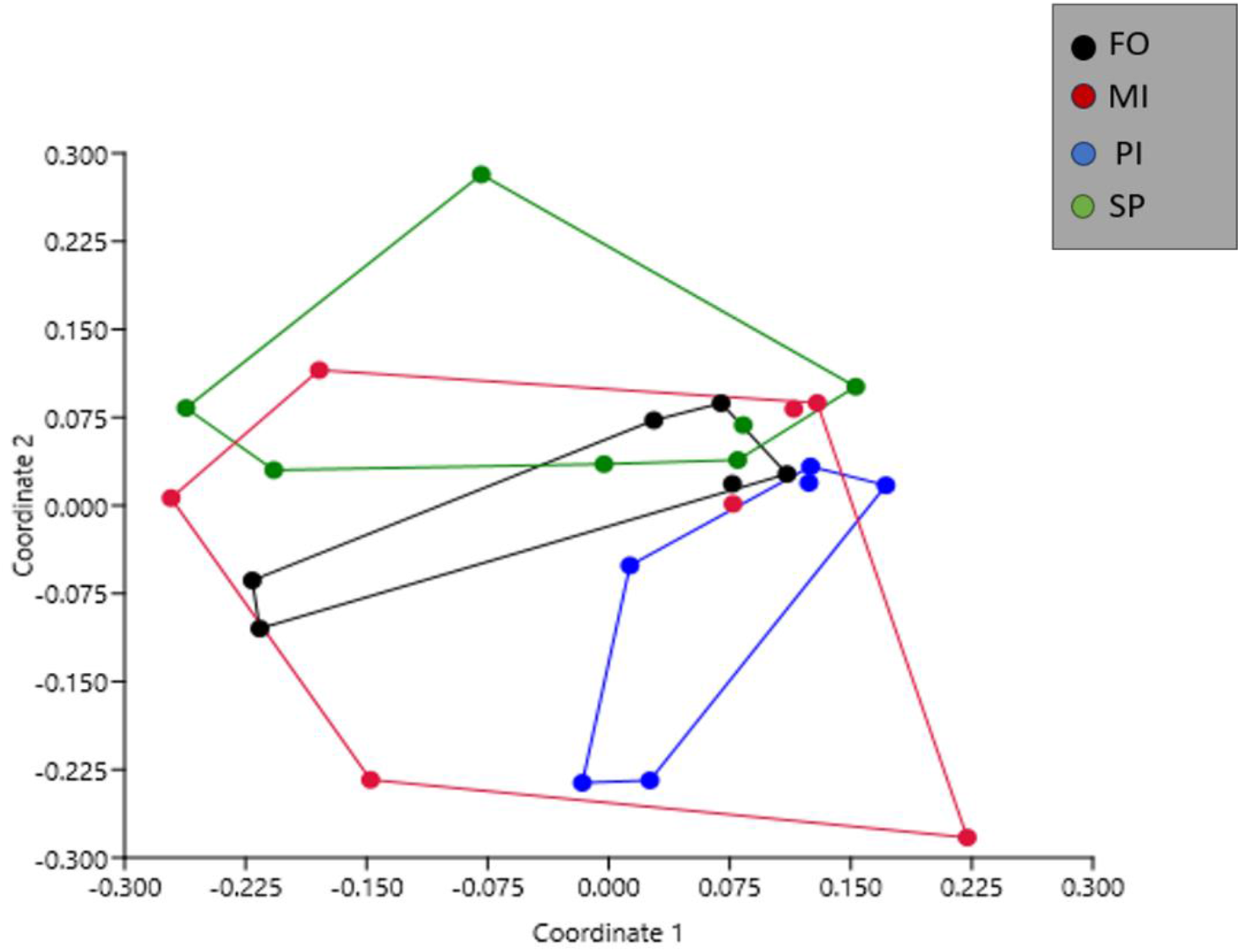
Non-metric multidimensional scaling (nMDS) plot for all the bacterial communities of all samples of the four dietary groups based on Bray-Curtis distances. PI samples are separated from FO and SP samples. FO: Fish Oil, MI: *Microchloropsis + Isochrysis*, SP: *Schizochytrium + Phaeodactylum*, PI: *Phaeodactylum* + *Isochrysis*.

The unique OTUs in FO, PI, SP and MI groups (Figure S1a) were 40 (7.7% of all OTUs), 61 (11.8%), 148 (28.5%) and 56 (10.8%) respectively. In total, 11.2% of the OTUs were found in all groups (FO, PI, SP, MI). Furthermore, there were 91 unique OTUs in FO compared to PI (30%), 95 unique OTUs compared to MI (34.2%) and 76 unique OTUs compared to SP (19.5%) (Figure S1b). Shared OTUs were in a higher proportion between FO and MI (34.5%) with FO and PI being very close (33%) and, lastly, FO and SP (29%). SP had the highest number of unique OTUs (199 OTUs - 51%) compared to FO, with MI and PI having smaller contributions of unique OTUs (87 OTUs - 31.3% and 112 OTUs -37%, respectively).

### Midgut bacterial taxonomic composition

Proteobacteria and Actinobacteriota were within the three most abundant phyla in all dietary groups (Figure S2). In FO and MI, the third most abundant phylum was Firmicutes, while in PI and SP the third most abundant phylum was Verrucomicrobiota and Bacteroidota, respectively. In FO, MI and SP, despite differences in either the abundance rank of the phyla (FO-MI) or the phyla presence (FO-SP), the ratios of the phyla abundances were quite close (Figure S2). In contrast, in PI there was a large increase in the relative abundance of Proteobacteria, since Proteobacteria were x2.1 more abundant than Actinobacteriota and x8.1 than Verrucomicrobiota. Actinobacteriota also had a large difference in relative abundance with the third most abundant phylum, Verrucomicrobiota (x3.8). Regarding other phyla, only in MI the Fusobacteriota occurred with a relative abundance of 4.3% and in PI and SP the Firmicutes with relative abundance of 5.9% and 13.9%, respectively.

In all feeding groups, the Mycobacteriaceae was the most abundant family in the FO (27.6% of reads), MI (28.4% of reads) and SP (27.2% of reads) and second most abundant in PI with relative abundance 22.0% of reads (Figure 2). In the PI treatment, the most abundant family was the Rhodobacteraceae (31.7% of reads), which in MI and SP occurred in very low abundances (2.9% and 1.3% of reads, respectively). These abundances were due to the most dominant OTUs, which in FO, MI and SP was OTU001 (27.6%, 28.3%, 27.1% of reads respectively) and OTU004 in PI with relative abundance 23.9% (Table S3). Nucleotide BLAST search assigned OTU001 as *Mycolicibacterium aurum* and OTU004 as *Yoonia litorea*. Compared to FO, in the other feeding groups the family Rhizobiaceae was not present and the family Xanthomonadaceae was only present in MI (<4% relative abundance). The family Vibrionaceae occurred in MI (8.3% of reads) and PI (12.3% of reads) but it was absent in SP and FO. Treatment MI had the highest number of families contributing >4% relative abundance (nine families), while SP had the lowest (five families).

**Figure 2.**
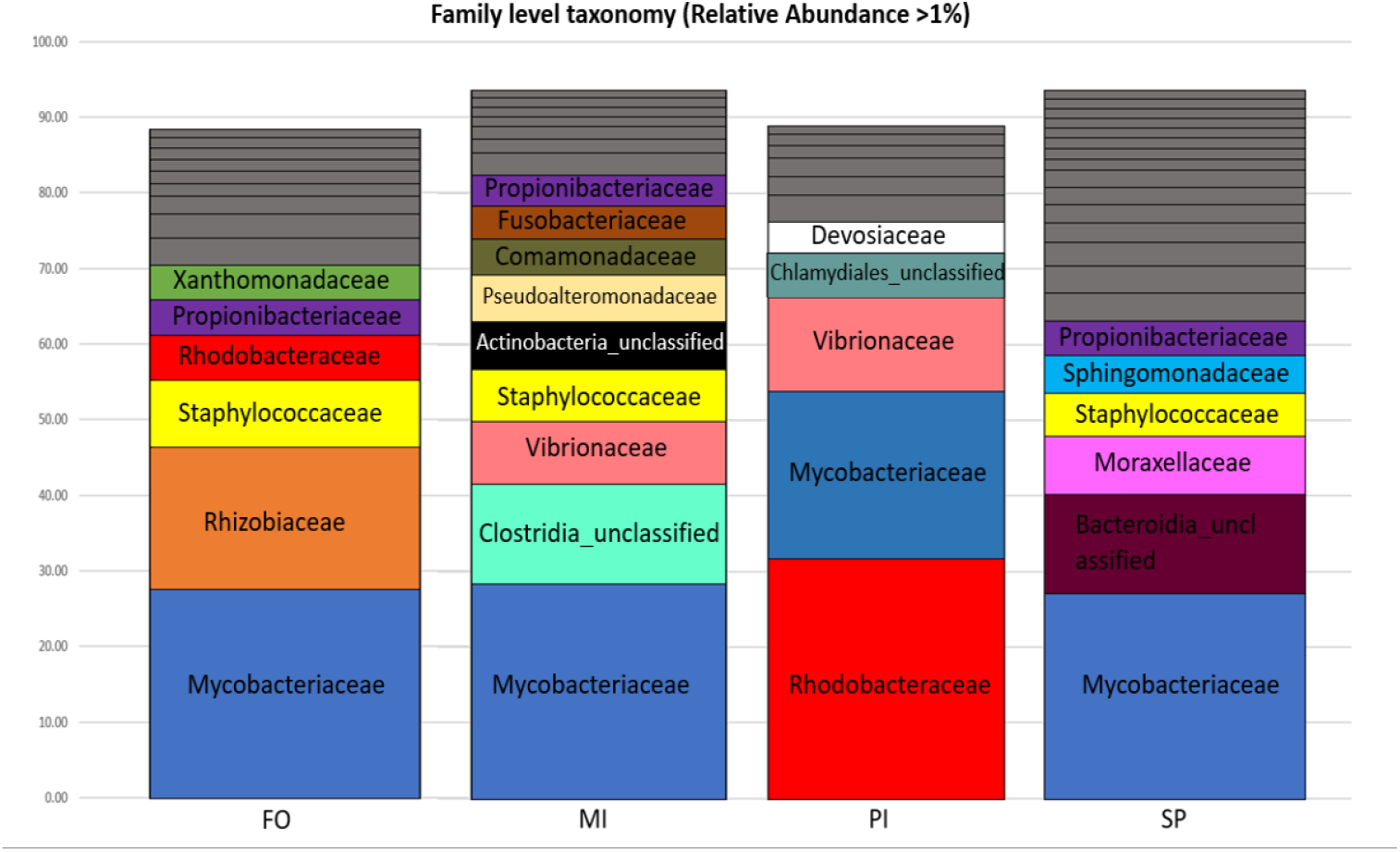
Taxonomic composition of the bacterial families in each dietary group. Relative abundances were calculated adding the relative abundance (calculated based on the average samples reads of the seven samples per diet) of all OTUs belonging to each family. Only the families with relative abundance > 1% were considered for the diagram. Families with relative abundance < 4% are depicted in grey. FO: Fish Oil, MI: *Microchloropsis + Isochrysis*, SP: *Schizochytrium + Phaeodactylum*, PI: *Phaeodactylum* + *Isochrysis*.

### Most important OTUs

In the dominant OTUs (cumulative relative dominance > 80% - Figure 3) three OTUs occurred in all feeding groups (OTU001, OTU003, OTU012).

**Figure 3.**
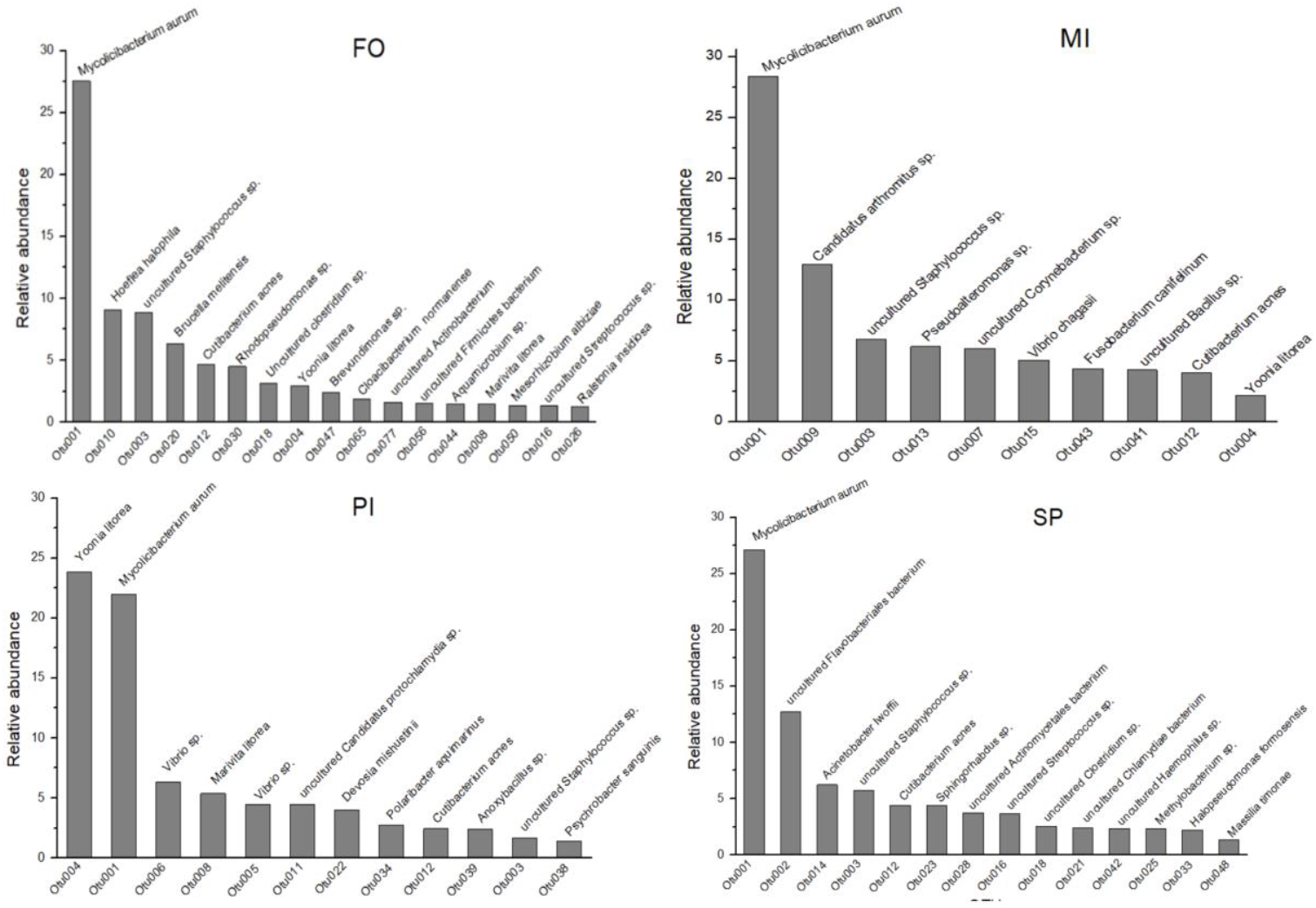
Most dominant OTUs in each dietary group (cumulative relative dominance > 80% and closest relative (Nucleotide BLAST) at the species level. Relative abundances are % of reads. FO: Fish Oil, MI: *Microchloropsis + Isochrysis*, SP: *Schizochytrium + Phaeodactylum*, PI: *Phaeodactylum* + *Isochrysis*.

OTU003 and OTU012 occurred in small proportions in PI (1.67% and 2.44% of reads, respectively). From the Nucleotide BLAST search, the OTU003 closest relative was the uncultured *Staphylococcus sp*. and OTU012 closest relative was *Cutibacterium acnes*. OTU004, which was the most dominant in PI, was among the most dominants of FO and MI, but with low relative abundances (2.95% and 2.16% of reads, respectively). Within the dominant OTUs there were several unique ones that occurred in each feeding group.

The qualitative differences already observed in phyla and families were also observed at the level of dominant OTUs in each group. To explore this further, we compared key OTUs between the FO and the three experimental groups. This comparison unveiled which OTUs occurred in high abundance and were common in both the FO and experimental feeds, or exclusive to the FO (Figure 4, Table S4). 11 common OTUs were highly abundant in FO compared to MI, PI and SP. Furthermore, among the highly abundant OTUs, two of them (OTU001 and OTU003) were observed in all treatments. Most of the highly abundant OTUs in MI, PI and SP were unique to each experimental feed, with only two OTUs (OTU006, OTU027) being common to FO-MI and FO-PI.

**Figure 4.**
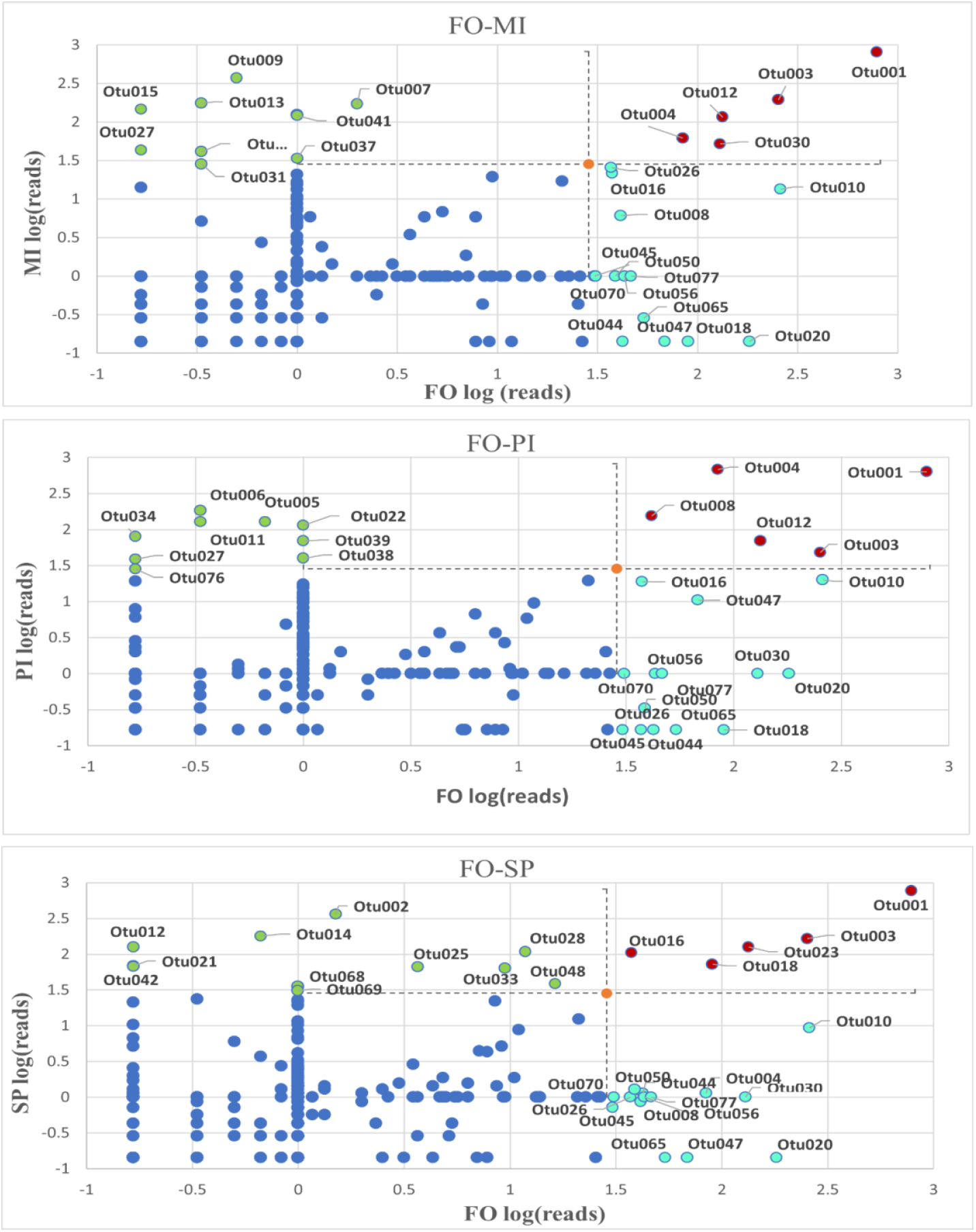
Relative abundance of the FO reads versus their respective values of the PI, MI and SP. OTUs in green dots are in ≥1% relative abundance in PI or MI or SP but not in FO, OTUs in red dots are in ≥1% relative abundance in FO and PI, MI and SP concomitantly, OTUs in pale blue dots are in ≥1% relative abundance in FO but not in PI, MI and SP and OTUs in dark blue dots are in ≤1% relative abundance in any treatment. The limit set (orange point) is the log of 1% of the total reads. FO: Fish Oil, MI: *Microchloropsis + Isochrysis*, SP: *Schizochytrium + Phaeodactylum*, PI: *Phaeodactylum* + *Isochrysis*.

### Presumptive functional pathways

Compared to FO, one metabolic pathway was considerably overexpressed (FUCCAT-PWY with values: LOG(tax.function.abundance PI/FO) = 1.988, LOG(tax.function.abundance MI/FO) = 1.988, LOG(tax.function.abundance SP/FO) = 1.536) and one considerably underexpressed (PWY-6470 with values: LOG(tax.function.abundance PI/FO) = -1.511, LOG(tax.function.abundance MI/FO) = - 1.618,, LOG(tax.function.abundance SP/FO) = -1.509) in all three experimental feeding groups (Figure 5, Table S5). According to the MetaCyc database, the FUCCAT-PWY pathway corresponds to L-fucose degradation, and the PWY-6470 pathway corresponds to peptidoglycan biosynthesis. Several common pathways were overexpressed in MI and PI relatively to FO and the highest number of unique pathways were overexpressed in SP. From the KEGG database it was found that the genomes of most of the OTUs in each of the three experimental feeds (see Fig. 4) contained all the genes encoding for the enzymes of the L-Fucose degradation pathway (Table S6).

**Figure 5.**
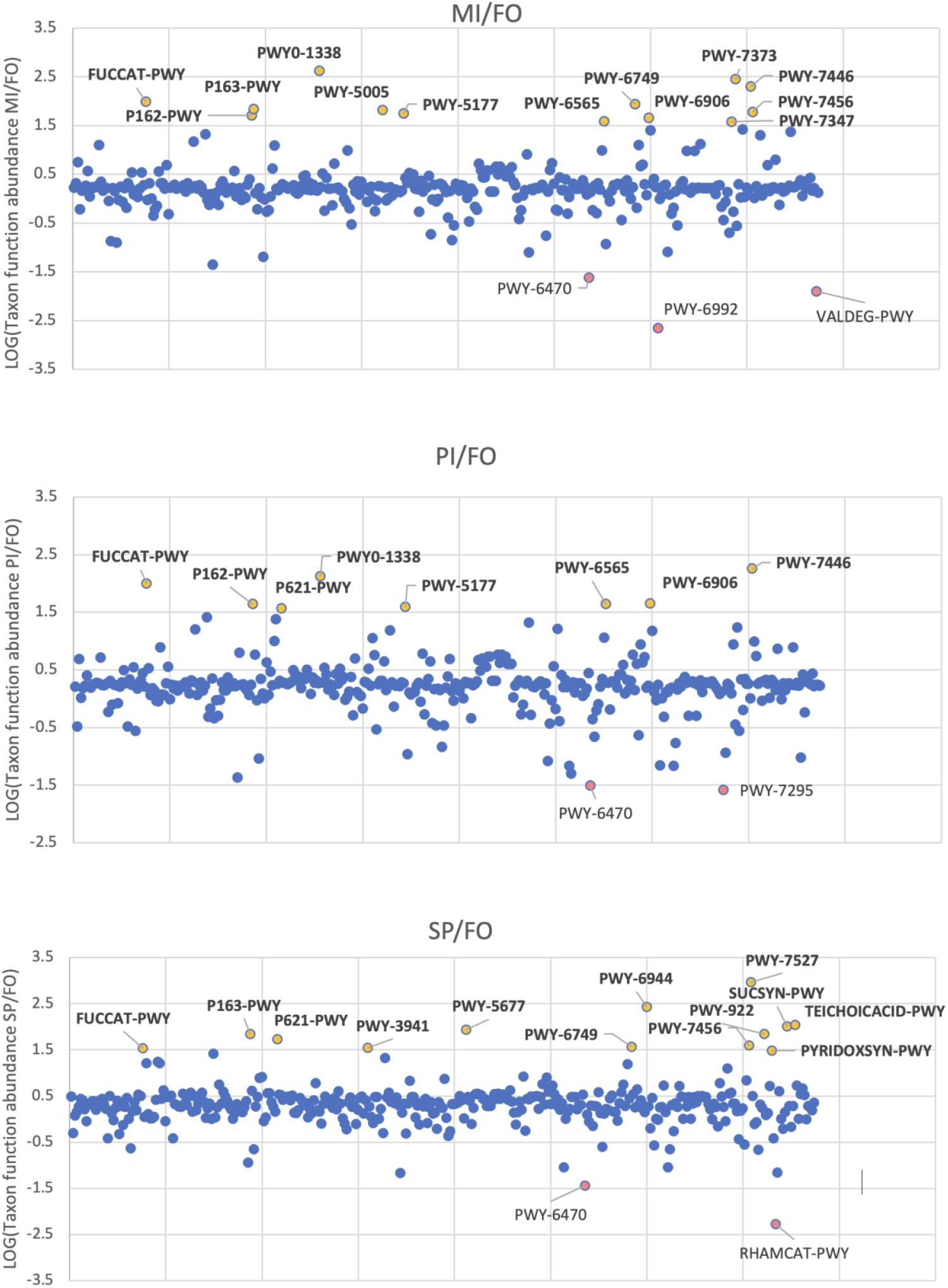
Scatter plots with every pathway in x axis and the log of the ratio PI/FO, MI/FO and SP/FO of the Taxon function abundance in y axis. Pathways > 1.5 are considered overexpressed in PI, MI and SP respectively compared to FO and pathways < -1.5 are considered underexpressed. FO: Fish Oil, MI: *Microchloropsis + Isochrysis*, SP: *Schizochytrium + Phaeodactylum*, PI: *Phaeodactylum* + *Isochrysis*.

## DISCUSSION

In this study, it was hypothesized that the inclusion of microalgal-based diets in experimental reared gilthead sea bream would induce structural bacterial microbiota changes adapted to these novel feed ingredients. In similar studies with the use of microalgae in the diet of gilthead sea bream, variable results have been reported. Cerezuela et al. (2012) reported that the dietary inclusion of 100g/kg *Tetraselmis chui* or *P. tricornutum* decreased the intestinal microbial richness with a lower Shannon index value than the control. In the current study, the Shannon index decreased in fish fed with the *Schizochytrium* - *P. tricornutum* (SP) diet but slightly increased in fish fed with the *P. tricornutum* – *Isochrysis* diet. Nevertheless, these two diets had more OTUs (Taxa_S) than the control (FO). Jorge et al. (2019) using *M. gaditana* for fishmeal replacement reported a higher, but not statistically significant different to the control, intestinal richness and diversity of the microbiota. Using a microalgae blend of *Tisochrysis lutea* (member of the *Isochrysis* spp. group), *M. gaditana* and *Scenedesmus almeriensis* at 5%, 15% and 25% dietary inclusion levels, Garcia-Marquez et al. (2023) found that the microalgae blend induced an increase in bacterial species diversity and a distinct shift in microbiota fingerprinting as inclusion levels increased. In our study, compared to the control (FO), the diet containing *M. gaditana* (MI) showed lower alpha diversity indices, but this was not statistically significant. Similar lack of statistically significance has been reported in the *Sparus aurata* gut microbiota after the inclusion of 5% hydrolysed *M. gaditana* (Cerezo-Ortega et al. 2021).

Studies with dietary inclusion of *Schizochytrium* sp. in Nile tilapia (Souza et al. 2020), zebrafish (Shi et al. 2021) and rainbow trout (Lyons et al. 2016) also showed varying results. Specifically, in Nile tilapia a diet with 1.2% inclusion of *Schizochytrium sp*. led to higher Inverse Simpson and Shannon values in the *Schizochytrium*-fed fish, but these were not significantly different to those in the control group. In zebrafish, the dietary inclusion at 0, 60 and 120g/kg of *Schizochytrium sp*. resulted in insignificant differences in Shannon index among the three groups, but in the beta diversity nMDS showed that both the *Schizochytrium sp*. supplemented diets were separated from the control group (Shi et al. 2021). The diet containing *Schizochytrium sp*. in our study (SP) had lower values in Shannon-H and Simpson_1-D indices than the control (FO), but these were not statistically significant and also there were no statistically significant differences in beta diversity compared to the control. Furthermore, a 5% inclusion of *Schizochytrium limacinum* for fish oil replacement in the diet of rainbow trout resulted in a greater level of microbial diversity (Lyons et al. 2016; a fact that was also observed in our results using the SP diet that had a higher OTUs (Taxa_S) index number than the control (FO).

Overall, the effect on gut microbiota diversity is primarily determined by the different species of microalgae, the fish as the host and the levels of inclusion of each microalga in the diet. The literature as discussed above in combination with the results of our study shows the various changes that occur as the gut microbiome adapts to the different components added by microalgae to the diet. However, differences in diversity alone cannot show the overall effects and it is necessary to investigate the structural changes at the level of bacterial phyla, families, and specific OTUs. As it has been shown that the phycosphere of cultivated microalgae used as feed might affect the fish gut microbiota in early developmental stages (e.g. Nikouli et al. 2019), in the future the impact of the bacteria contained in the microalgae to be incorporated in the aquafeed should be investigated as well.

### Most important OTUs

The dominant phyla found (Proteobacteria, Actinobacteriota, Bacteroidota) are common in the gilthead seabream gut microbiome (Kormas et al. 2014, Nikouli et al. 2018, 2019, Cerezo-Ortega et al 2021) and also in other fish species gut microbiome (Llewellyn et al. 2016, Rosenau et al. 2021, Zhang et al. 2022). In PI, however, there was an increased occurrence of Verrucomicrobiota. The Mycobacteriaceae family is the one that is highly abundant in all groups, due to OTU001, which belongs to the genus *Mycolicibacterium*. From the available literature it seems that this genus is not common in the gilthead sea bream gut. It includes some species of pathogenic micro-organisms for humans (*Mycobacterium tuberculosis, Mycobacterium leprae*) but also several species that inhabit a diverse range of environments including water bodies, soil, and metal-working fluids (Gupta, Lo & Son 2018). *Mycobacterium* genus has also been found in the gut of freshwater species (Gallet et al. 2022). The closest relative, *M. aurum* has never been reported as a pathogen, but its possible functions in the gut remain unknown. The Rhodobacteraceae family member OTU004 is the dominant in the PI treatment, closely related to *Yoonia litorea* (Paracocacceae family). *Y. litorea* though has been recently classified in the *Roseobacter* clade within the Rhodobacteraceae family (Wirth & Whitman 2018). Later, it was proposed the family Roseobacteraceae for members of the *Roseobacter* clade (Liang et al. 2021). Misclassifications have been a recurring problem between the Paracocacceae and Roseobacteraceae families (Zhang et al. 2023), but *Yoonia* sp. isolates are phylogenetically placed in the Roseobacteraceae family (Feng & Xing 2023). Species from the *Roseobacter* clade have been isolated from seawater marine sediments and biofilms, and are often associated with phytoplankton, macroalgae and marine animals (Wirth & Whitman 2018). They have also been found in intestine of finfish and they seem to have probiotic effects (Ringø et al. 2022). Specifically, for the *Yoonia-Loktanella* group it has been speculated that they offer defensive secondary metabolites to deep-sea sponges (Lo Giudice et al. 2024).

OTU003 was highly abundant in FO and MI, PI and SP and its closest relative is *Staphylococcus* sp., while OTU012 was highly abundant in FO, MI and PI and closely related to *Cutibacterium acnes. Staphylococcus* sp. is a common bacterial genus in healthy fish gut microbiomes and some species may have probiotic effects (Zorriehzahra et al.2015, Borah et al. 2016, El-Saadony et al. 2021). *Cutibacterium acnes* has been found in other fish species and it has been suggested to have potential probiotic features (Nikouli et al. 2021). OTU006 and OTU027, that belong to the *Vibrio* genus, were in high abundance in MI and PI compared to FO. *Vibrio* is a common bacterial genus found in fish gut with both pathogenic and non-pathogenic species. Some *Vibrio* species can act as symbionts assisting in the breakdown of complex dietary components, such as polysaccharides, with strains found to produce amylase, cellulose and chitinase among others (Egerton et al. 2018). From the rest top abundant OTUs, in MI OTU013, OTU031 and OTU041 were closely related to *Pseudoalteromonas* sp., *Pseudomonas* sp. and *Bacillus* sp. respectively. Several species of these genera are known to have probiotic effects in the fish gut (Mladineo et al. 2016, Zorriehzahra et al. 2015, Salam et al. 2021). OTU030 was abundant in FO and MI and was related to *Rhodopseudomonas* sp., a genus that is also reported to have probiotic effects in the fish gut (Koga et al. 2022). Thus, from the point of structural changes of the bacterial communities, the MI experimental feed seems to promote several beneficial bacterial species in the gut of gilthead sea bream.

L-fucose degradation was the only considerably overexpressed bacterial metabolic pathway in the three experimental feeds in comparison to FO. Fucose is a deoxyhexose that is found as a part of the polysaccharide content of microalgae (Bernaerts et al. 2018, Wan et al. 2019). It is also commonly found in animal cell glycans, macroalgae, cyanobacteria and microbial exopolysaccharides (Becker & Lowe 2003). Furthermore, search of the KEGG database confirmed that most of the important bacterial genera found are affiliated to genomes with all five enzymes to carry out the L-fucose degradation. This suggests that the microalgal-based diets induced a clear gut bacterial response, at the functional level, favoring the proliferation of bacteria which benefit from specific microalgal compounds with potential benefits to the host, too. The degradation of fucose is a likely beneficial-to-the-host service. For example, in humans fucose degraders are known to induce the production of the beneficial short-chain fatty acids (Morrison et al. 2016, Høgsgaard et al. 2023, Fusco et al. 2023). The metabolic fate of the fucose degradation and its possible effect on the sea bream’s physiology remains to be specifically investigated in the future.

The only pathway that was considerably under-expressed in the three experimental feeds compared to FO is peptidoglycan biosynthesis. The cell wall of some microalgae species contains peptidoglycan (Agboola et al. 2019, Machado et al. 2022), but the species used in our research do not seem to contain it (Domozych et al. 2012, Le Costaouëc et al. 2017, Nadzir et al. 2023). Peptidoglycan forms around 90% of the dry weight of Gram-positive bacteria but only 10% of Gram-negative strains (Malanovic & Lohner 2016). The relative abundance of Gram-negative phyla and families like Vibrionaceae in PI and MI and Bacteroidota in SP is higher than FO. Consequently, the putative under-expression of peptidoglycan synthesis with the three diets compared to FO is likely representative of the higher presence of Gram-taxa. Finally, SP was the only treatment with unique overexpressed pathways along with the highest number of unique OTUs compared to FO for which, however, only one of the treatment’s unique OTUs was included in these overexpressed pathways (Table S7). This microalgal feed inclusion requires further investigation with in vitro experiments using specific isolates and growth media to test the functionality of these presumptive metabolic pathways.

In conclusion, this study showed that the inclusion of three microalgal-based diets induced both structural and functional changes in the sea bream’s gut microbiome. The structural changes in each dietary treatment were considerable compared to the control diet. The putative under-expression of peptidoglycan synthesis with the 3 diets compared to FO was likely representative of the higher presence of Gram-strains. However, the total midgut bacterial communities which were favoured by the microalgal inclusion were inferred to be able to metabolise L-fucose, a major microalgal carbohydrate.

## Supporting information

Supplementary material contains two tables and two figures

## AKNOWLEDGMENTS

This work was financially supported by the English MSc Program “Host-Microbe Interactions of the University of Thessaly and co-financed by the European Union and Greek co-funded project entitled “Use of insect protein and microalgal oil for fishmeal and fish oil replacement in the diets of gilthead seabream (*Sparus aurata*) and European seabass (*Dicentrarchus labrax*) under national funds through the Operational Programme “Competitiveness, Entrepreneurship and Innovation-EPAnEK 2014-2020” (project code: MIS 5045804)).

