## Supplementary material contains two tables and two figures for "The effect of dietary fish oil replacement by microalgae on the gilthead sea bream midgut bacterial microbiota"

**Table S1.** PERMANOVA pairwise test for alpha diversity metrics based on Bray – Curtis distance matrix. Statistically significant differences for p < 0.05 are shown in red. FO: Fish Oil, PI: *Phaeodactylum* + *Isochrysis*, SP: *Schizochytrium + Phaeodactylum*, ΜΙ: *Microchloropsis + Isochrysis.*

| **Taxa_S** | | | | |
| --- | --- | --- | --- | --- |
| Aquafeed | FO | PI | SP | MI |
| FO | - | 0.636 | 0.411 | 0.106 |
| PI |  | - | 0.342 | 0.046 |
| SP |  |  | - | 0.718 |
| MI |  |  |  | - |
| **Chao-1** | | | | |
| Aquafeed | FO | PI | SP | MI |
| FO | - | 0.429 | 0.547 | 0.220 |
| PI |  | - | 0.562 | 0.442 |
| SP |  |  | - | 0.623 |
| MI |  |  |  | - |
| **Shannon-H** | | | | |
| Aquafeed | FO | PI | SP | MI |
| FO | - | 0.911 | 0.470 | 0.124 |
| PI |  | - | 0.286 | 0.049 |
| SP |  |  | - | 0.529 |
| MI |  |  |  | - |
| **Simpson 1-D** | | | | |
| Aquafeed | FO | PI | SP | MI |
| FO | - | 0.940 | 0.412 | 0.164 |
| PI |  | - | 0.388 | 0.112 |
| SP |  |  | - | 0.632 |
| MI |  |  |  | - |

**Table S2**. PERMANOVA pairwise test for all samples based on Bray – Curtis distance matrix. Statistically significant differences for p<0.05 are shown in red. FO: Fish Oil, PI: *Phaeodactylum* + *Isochrysis*, SP: *Schizochytrium + Phaeodactylum*, ΜΙ: *Microchloropsis + Isochrysis.*

| Aquafeed | FO | PI | SP | MI |
| --- | --- | --- | --- | --- |
| FO | - | 0.034 | 0.817 | 0.790 |
| PI |  | - | 0.024 | 0.089 |
| SP |  |  | - | 0.851 |
| MI |  |  |  | - |

**Table S3.** Most abundant OTUs in each feeding group. FO: Fish Oil, ΜΙ: *Microchloropsis + Isochrysis*, SP: *Schizochytrium + Phaeodactylum*, PI: *Phaeodactylum* + *Isochrysis*.

| **Aquafeed** | **Relative Abundance of the most abundant OTU (% of reads), closest relative (>97%)** | **No. of dominant OTUs with ≥80% cumulative relative abundance** |
| --- | --- | --- |
| **FO** | OTU-0001 (27.6%), *Mycolicibacterium aurum* | 17 |
| **MI** | OTU-0001 (28.3%), *Mycolicibacterium aurum* | 10 |
| **SP** | OTU-0001 (27.1%), *Mycolicibacterium aurum* | 14 |
| **PI** | OTU-0004 (23.9%), *Yoonia litorea* | 12 |

**Table S4.** Taxonomy and closest relative of the OTUs with the GenBank accession numbers (Nucleotide BLAST) depicted in Figure 7. FO: Fish Oil, ΜΙ: *Microchloropsis + Isochrysis*, SP: *Schizochytrium + Phaeodactylum*, PI: *Phaeodactylum* + *Isochrysis*.

| FO high abundance OTUs compared to the three experimental feed groups | | | |
| --- | --- | --- | --- |
| 11 common OTUs in FO-PI, FO-MI and FO-SP | | | |
| OTU010 | Proteobacteria | Rhizobiaceae | *Hoeflea halophila (100.00%)* - MT628728.1 |
| OTU020 | Proteobacteria | Rhizobiaceae | *Brucella melitensis (99.25%)* - MT611105.1 |
| OTU026 | Proteobacteria | Burkholderiales_unclassified | *Ralstonia insidiosa (99.53%)* - MK968569.1 |
| OTU044 | Proteobacteria | Rhizobiaceae | *Aquamicrobium sp. (98.26%)* - MT457451.1 |
| OTU045 | Proteobacteria | Coxiellaceae | *Coxiella-like endosymbiont (96.04%)* - MH645185.1 |
| OTU047 | Proteobacteria | Caulobacteraceae | *Brevundimonas sp. (100.00%)* - MN620407.1 |
| OTU050 | Proteobacteria | Rhizobiaceae | *Mesorhizobium albiziae (100.00%)* - KP337606.1 |
| OTU056 | Acidobacteriota | Blastocatellaceae | uncultured *Firmicutes bacterium (100.00%)* - EU298762.1 |
| OTU065 | Bacteroidota | Weeksellaceae | *Cloacibacterium normanense (100.00%)* - MK294295.1 |
| OTU070 | Bacteroidota | Weeksellaceae | *Chryseobacterium sp. (100.00%)* - CP050993.1 |
| OTU077 | Actinobacteriota | Actinobacteriota_unclassified | *uncultured Actinobacterium (97.66%)* - JN178908.1 |
| 2 common OTUs in FO-PI and FO-MI | | | |
| OTU016 | Firmicutes | Streptococcaceae | *uncultured Streptococcus sp. (100.00%)* - LT681499.1 |
| OTU018 | Firmicutes | Clostridiaceae | *Uncultured clostridium sp. (99.75%)* - MN663533.1 |
| 1 common OTU in FO-PI and FO-SP | | | |
| OTU030 | Proteobacteria | Xanthobacteraceae | *Rhodopseudomonas sp. (100.00%)* - MN555639.1 |
| 1 common OTU in FO-MI and FO-SP | | | |
| OTU008 | Proteobacteria | Rhodobacteraceae | *Marivita litorea (99.25%)* - MN746209.1 |
| 1 OTU included exclusively in FO-SP | | | |
| OTU004 | Proteobacteria | Rhodobacteraceae | *Yoonia litorea (99.25%)* - MZ443920.1 |
| Common high abundance OTUs of FO with MI, PI and SP | | | |
| 2 common OTUs in FO-PI, FO-MI and FO-SP | | | |
| OTU001 | Actinobacteriota | Mycobacteriaceae | *Mycolicibacterium aurum (99.26%)* -MT406679.1 |
| OTU003 | Firmicutes | Staphylococcaceae | uncultured *Staphylococcus sp. (99.77%)* - FJ957685.1 |
| 2 common OTUs in FO-PI and FO-MI | | | |
| OTU004 | Proteobacteria | Rhodobacteraceae | *Yoonia litorea (99.25%)* - MZ443920.1 |
| OTU012 | Actinobacteriota | Propionibacteriaceae | *Cutibacterium acnes (100.00%)* - MT613579.1 |
| 1 OTU included exclusively in FO-PI | | | |
| OTU008 | Proteobacteria | Rhodobacteraceae | *Marivita litorea (99.25%)* -MN746209.1 |
| 1 OTU included exclusively in FO-MI | | | |
| OTU030 | Proteobacteria | Xanthobacteraceae | *Rhodopseudomonas sp. (100.00%)* - MN555639.1 |
| 3 OTUs included exclusively in FO-SP | | | |
| OTU016 | Firmicutes | Streptococcaceae | *uncultured Streptococcus sp. (100.00%)* - LT681499.1 |
| OTU018 | Firmicutes | Clostridiaceae | *Uncultured clostridium sp. (99.75%)* - MN663533.1 |
| OTU023 | Proteobacteria | Sphingomonadaceae | *Sphingorhabdus sp. (99.75%)* -MK493524.1 |
| ΜΙ, PI and SP high abundance OTUs compared to FO | | | |
| 2 common OTUs in FO-PI and FO-MI | | | |
| OTU006 | Proteobacteria | Vibrionaceae | *Vibrio sp. (99.53%)* - KY914442.1 |
| OTU027 | Proteobacteria | Vibrionaceae | *Vibrio sp. (99.77%)* - MT634723.1 |
| 7 OTUs included exclusively in FO-PI | | | |
| OTU005 | Proteobacteria | Vibrionaceae | *Vibrio sp. (99.30*%) - MN252252.1 |
| OTU011 | Verrucomicrobiota | Chlamydiales_unclassified | uncultured *Candidatus protochlamydia sp. (91.82%)* - OP914912.1 |
| OTU022 | Proteobacteria | Devosiaceae | *Devosia mishustinii (98.49%)* - KX982774.1 |
| OTU034 | Bacteroidota | Flavobacteriaceae | *Polaribacter aquimarinus (100.00%)* - OP344420.1 |
| OTU038 | Proteobacteria | Moraxellaceae | *Psychrobacter sanguinis (99.30%)* -MN758810.1 |
| OTU039 | Firmicutes | Bacillaceae | *Anoxybacillujs sp. (100.00%)* -MN885758.1 |
| OTU076 | Proteobacteria | Rhodobacteraceae | *Planktotalea sp. (99.75%*) - KP172216.1 |
| 7 OTUs included exclusively in FO-MI | | | |
| OTU007 | Actinobacteriota | Actinobacteria_unclassified | uncultured *Corynebacterium sp. (99.76%)* - LT692791.1 |
| OTU009 | Firmicutes | Clostridia_unclassified | *Candidatus arthromitus sp. (98.01%)* - AP012210.1 |
| OTU013 | Proteobacteria | Pseudoalteromonadaceae | *Pseudoalteromonas sp. (99.53%)* - MF975560.1 |
| OTU015 | Proteobacteria | Vibrionaceae | *Vibrio chagasii (99.30%)* - MT269630.1 |
| OTU031 | Proteobacteria | Pseudomonadaceae | *Pseudomonas sp. (99.30%)* - MT585910.1 |
| OTU037 | Verrucomicrobiota | Chlamydiales_unclassified | *Uncultured Chlamydiales bacterium (96.96%)* - KC902452.1 |
| OTU041 | Proteobacteria | Comamonadaceae | *uncultured Bacillus sp. (99.53%)* - EU567044.2 |
| 11 OTUs included exclusively in FO-SP | | | |
| OTU002 | Bacteroidota | Bacteroidia_unclassified | *uncultured Flavobacteriales bacterium (91.51%)* - FJ425622.1 |
| OTU012 | Actinobacteriota | Propionibacteriaceae | *Cutibacterium acnes (100.00%)* - MT613579.1 |
| OTU014 | Proteobacteria | Moraxellaceae | *Acinetobacter Iwoffii (99.77%)* - MT626722.1 |
| OTU021 | Verrucomicrobiota | Simkaniaceae | *uncultured Chlamydiae bacterium (91.59%)* - DQ996921.1 |
| OTU025 | Proteobacteria | Beijerinckiaceae | *Methylobacterium sp. (99.25%)* -LC546537.1 |
| OTU028 | Actinobacteriota | Corynebacteriaceae | uncultured *Actinomycetales bacterium (99.26%)* - HM076482.1 |
| OTU033 | Proteobacteria | Pseudomonadaceae | *Halopseudomonas formosensis (99.06%)* - MT313134.1 |
| OTU042 | Proteobacteria | Pasteurellaceae | *uncultured Haemophilus sp. (100.00%)* - ON786180.1 |
| OTU048 | Proteobacteria | Oxalobacteraceae | *Massilia timonae (100.00%)* - MN932364.1 |
| OTU068 | Firmicutes | Lachnospiraceae | *Blautia sp. Marseille-P3313 (97.26%)* - LT631509.1 |
| OTU069 | Actinobacteriota | Actinobacteriota_unclassified | *Uncultured Solirubrobacterales bacterium (100.00%)* - JQ401013.1 |

**Table S5.** Overexpressed and underexpressed Pathways (from Metacyc database) in PI, MI and SP compared to FO. FO: Fish Oil, ΜΙ: *Microchloropsis + Isochrysis*, SP: *Schizochytrium + Phaeodactylum*, PI: *Phaeodactylum* + *Isochrysis*.

| **Overexpressed pathways in MI, PI, SP compared to FO** | |
| --- | --- |
| 1 common Pathway in PI/FO, MI/FO and SP/FO | |
| **FUCCAT-PWY** | **Degradation → L-Fucose Degradation** |
| 6 common Pathways in PI/FO and MI/FO | |
| PWY-7446 | Carbohydrate Degradation → Sulfoquinovose Degradation |
| PWY0-1338 | Lipopolysaccharide Biosynthesis |
| PWY-6906 | Carbohydrate Degradation → Sugar → Chitin |
| P162-PWY | Proteinogenic Amino Acid Degradation → L-glutamate Degradation |
| PWY-6565 | Amide, Amidine, Amine, and Polyamine Biosynthesis |
| PWY-5177 | Carboxylic Acid Degradation |
| 3 common Pathways in MI/FO and SP/FO | |
| PWY-6749 | Sugar Biosynthesis → Sugar → sialic acids |
| P163-PWY | Proteinogenic Amino Acid Degradation → L-glutamate Degradation |
| PWY-7456 | Carbohydrate Degradation → Polysaccharide Degradation→ plant mannan degradation |
| 1 common Pathway in PI/FO and SP/FO | |
| P621-PWY | caprolactam degradation |
| **Underexpressed Pathways in MI, PI, SP compared to FO** | |
| 1 common Pathway in PI/FO, MI/FO and SP/FO | |
| **PWY-6470** | **Cell Wall Biosynthesis → Peptidoglycan Biosynthesis** |
| 1 Pathway included exclusively in PI/FO | |
| PWY-7295 | Sugar Degradation → L-arabinose Degradation |
| 2 Pathways included exclusively in MI/FO | |
| PWY-6992 | Sugar Degradation → anhydrofructose pathway |
| VALDEG-PWY | Proteinogenic Amino Acid Degradation → L-valine Degradation |
| 1 Pathway included exclusively in SP/FO | |
| RHAMCAT-PWY | Sugar Degradation → L-rhamnose Degradation |

**Table S6.** The genus-level OTUs that contain all the enzymes of L-fucose degradation (from the KEGG database). Numbers indicate KEGG enzyme codes. + = it contains the enzyme, - = it does not contain the enzyme. Numbers indicate enzyme code.

| **OTUs** | **Genus** | **L-fucose mutarotase [5.1.3.29]** | **L-fucose isomerase [5.3.1.25]** | **L-fuculokinase [2.7.1.51]** | **L-fuculose-phosphate aldolase [4.1.2.17]** |
| --- | --- | --- | --- | --- | --- |
| OTU001 | *Mycolicibacterium* | + | + | + | + |
| OTU003 | *Staphylococcus* | + | + | + | + |
| OTU004 | *Yoonia* | + | + | + | + |
| OTU012 | *Cutibacterium* | + | + | + | + |
| OTU008 | *Marivita* | - | - | - | - |
| OTU030 | *Rhodopseudomonas* | + | + | + | + |
| OTU016 | *Streptococcus* | + | + | + | + |
| OTU018 | *Clostridium* | + | + | + | + |
| OTU023 | *Sphingorhabdus* | + | + | + | + |
| OTU005, 006, 027 | *Vibrio* | + | + | + | + |
| OTU011 | *Candidatus* Protochlamydia | + | + | + | + |
| OTU022 | *Devosia* | + | + | + | + |
| OTU034 | *Polaribacter* | + | + | + | + |
| OTU038 | *Psychrobacter* | + | + | + | + |
| OTU039 | *Ano+ybacillus* | + | + | + | + |
| OTU076 | *Planktotalea* | - | - | - | - |
| OTU007 | *Corynebacterium* | + | + | + | + |
| OTU009 | *Candidatus* Arthromitus | + | + | + | + |
| OTU013 | *Pseudoalteromonas* | + | + | + | + |
| OTU031 | *Pseudomonas* | + | + | + | + |
| OTU041 | *Bacillus* | + | + | + | + |
| OTU012 | *Cutibacterium* | + | + | + | + |
| OTU014 | *Acinetobacter* | + | + | + | + |
| OTU025 | *Methylobacterium* | + | + | + | + |
| OTU033 | *Halopseudomonas* | + | + | + | + |
| OTU042 | *Haemophilus* | + | + | + | + |
| OTU048 | *Massilia* | + | + | + | + |
| OTU068 | *Blautia* | + | + | + | + |

**Table S7**. Overexpressed SP pathways and OTUs with the higher taxon function abundance in each metabolic pathway. OTUs unique in the SP treatment are depicted with *

| **PWY-3941 - β-alanine biosynthesis II** | **Abundance** |
| --- | --- |
| OTU001 | 41.02 |
| OTU019 | 24.78 |
| OTU036 | 20.26 |
| OTU010 | 13.51 |
| OTU029 | 10.84 |
| **PWY-5677-succinate fermentation to butanoate** |  |
| OTU014 | 201.74 |
| OTU010 | 63.80 |
| OTU036 | 47.76 |
| OTU018 | 28.43 |
| **PWY-6944-androstenedione degradation I (aerobic)** |  |
| OTU001 | 3972.25 |
| OTU036 | 824.84 |
| OTU019 | 302.69 |
| OTU010 | 220.15 |
| OTU029 | 132.34 |
| **PWY-7527-L-methionine salvage cycle III** |  |
| OTU001 | 19.97 |
| OTU002 | 19.67 |
| OTU025 | 9.64 |
| **PWY-922 - mevalonate pathway I (eukaryotes and bacteria)** |  |
| OTU001 | 819.79 |
| OTU016 | 810.29 |
| *OTU069 | 171.64 |
| OTU036 | 148.06 |
| OTU025 | 137.70 |
| OTU010 | 111.16 |
| **PYRIDOXSYN-PWY - pyridoxal 5'-phosphate biosynthesis I** |  |
| OTU014 | 4203.82 |
| OTU001 | 3123.00 |
| OTU017 | 1349.27 |
| OTU019 | 1078.93 |
| OTU036 | 881.17 |
| **SUCSYN-PWY - sucrose biosynthesis I (from photosynthesis)** |  |
| OTU001 | 820.66 |
| OTU025 | 573.00 |
| OTU042 | 466.80 |
| OTU017 | 397.23 |
| OTU028 | 342.05 |
| **TEICHOICACID-PWY - poly(glycerol phosphate) wall teichoic acid biosynthesis** |  |
| OTU001 | 8.33 |
| OTU017 | 2.15 |
| OTU002 | 1.84 |

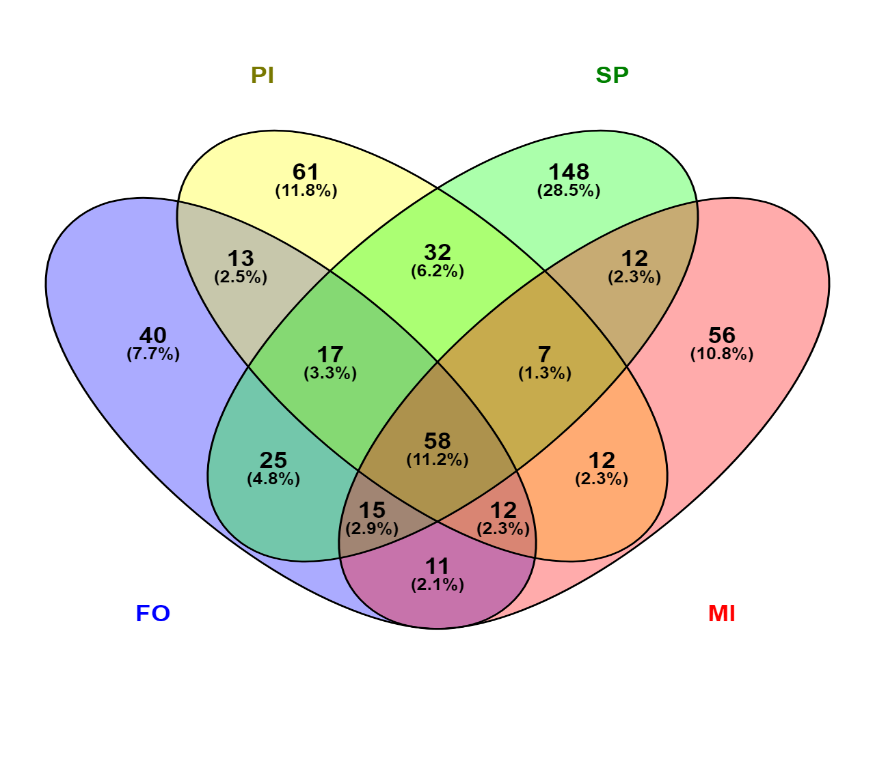

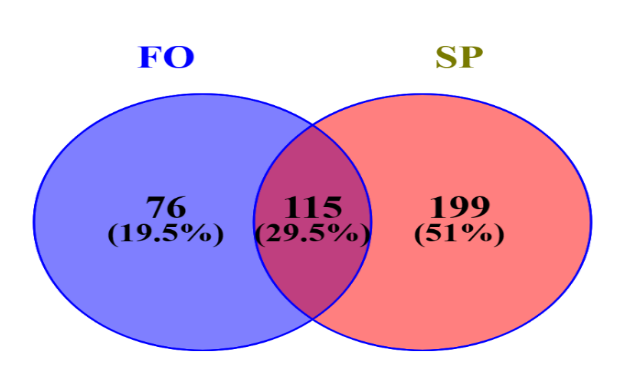

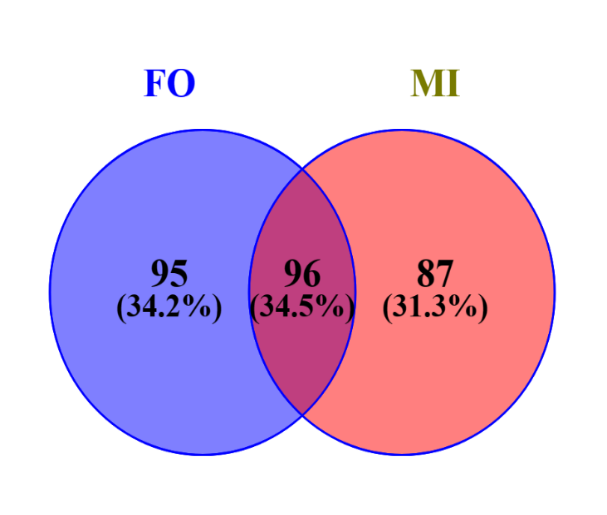

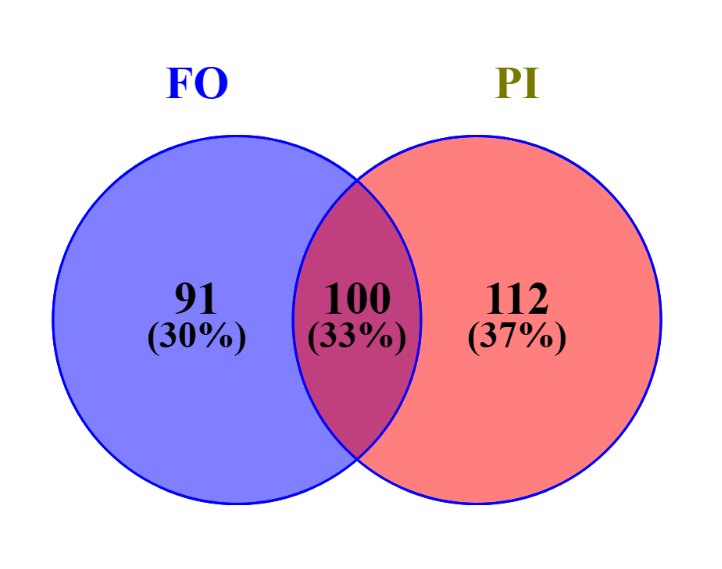

**Figure S1. (a)** Venn diagram demonstrating the number of common and unique OTUs and their percentages in the four feeding groups (FO, PI, SP, MI). (**b)** Venn diagrams demonstrating the numbers and percentages of the unique and common OTUs between the three experimental feed groups (MI, PI, SP) and the control (FO). FO: Fish Oil, ΜΙ: *Microchloropsis + Isochrysis*, SP: *Schizochytrium + Phaeodactylum*, PI: *Phaeodactylum* + *Isochrysis*.

a

bbb

**Fi**g**ure S2.** Relative Abundance ratio of the three most dominant phyla in each feeding group. Relative abundances were calculated from the average sample reads of each OTU. FO: Fish Oil, ΜΙ: *Microchloropsis + Isochrysis*, SP: *Schizochytrium + Phaeodactylum*, PI: *Phaeodactylum* + *Isochrysis*.

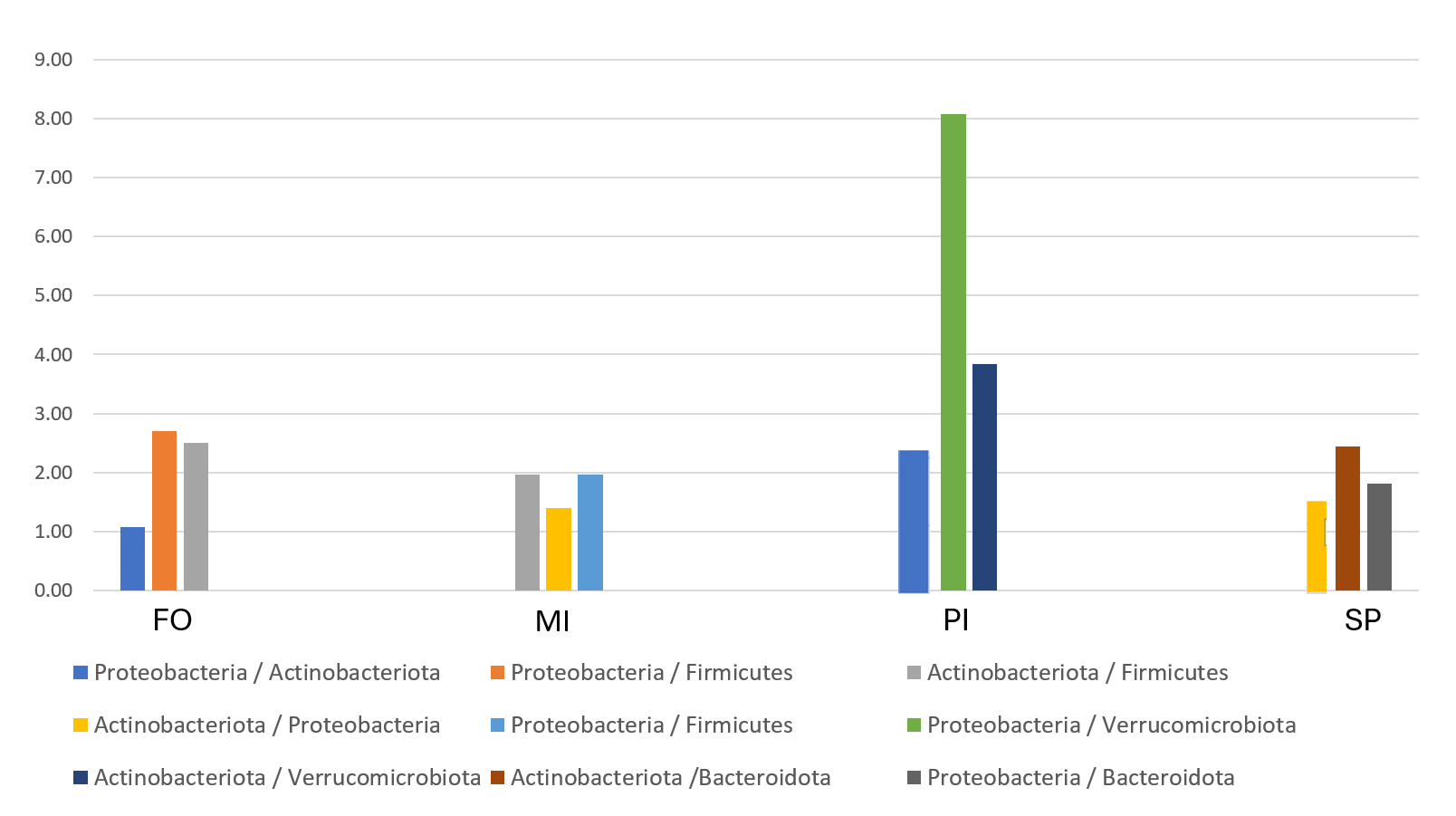
